# BRAND: A platform for closed-loop experiments with deep network models

**DOI:** 10.1101/2023.08.08.552473

**Authors:** Yahia H. Ali, Kevin Bodkin, Mattia Rigotti-Thompson, Kushant Patel, Nicholas S. Card, Bareesh Bhaduri, Samuel R. Nason-Tomaszewski, Domenick M. Mifsud, Xianda Hou, Claire Nicolas, Shane Allcroft, Leigh R. Hochberg, Nicholas Au Yong, Sergey D. Stavisky, Lee E. Miller, David M. Brandman, Chethan Pandarinath

## Abstract

Artificial neural networks (ANNs) are state-of-the-art tools for modeling and decoding neural activity, but deploying them in closed-loop experiments with tight timing constraints is challenging due to their limited support in existing real-time frameworks. Researchers need a platform that fully supports high-level languages for running ANNs (e.g., Python and Julia) while maintaining support for languages that are critical for low-latency data acquisition and processing (e.g., C and C++). To address these needs, we introduce the Backend for Realtime Asynchronous Neural Decoding (BRAND). BRAND comprises Linux processes, termed *nodes*, which communicate with each other in a *graph* via streams of data. Its asynchronous design allows for acquisition, control, and analysis to be executed in parallel on streams of data that may operate at different timescales. BRAND uses Redis to send data between nodes, which enables fast inter-process communication and supports 54 different programming languages. Thus, developers can easily deploy existing ANN models in BRAND with minimal implementation changes. In our tests, BRAND achieved <600 microsecond latency between processes when sending large quantities of data (1024 channels of 30 kHz neural data in 1-millisecond chunks). BRAND runs a brain-computer interface with a recurrent neural network (RNN) decoder with less than 8 milliseconds of latency from neural data input to decoder prediction. In a real-world demonstration of the system, participant T11 in the BrainGate2 clinical trial performed a standard cursor control task, in which 30 kHz signal processing, RNN decoding, task control, and graphics were all executed in BRAND. This system also supports real-time inference with complex latent variable models like Latent Factor Analysis via Dynamical Systems. By providing a framework that is fast, modular, and language-agnostic, BRAND lowers the barriers to integrating the latest tools in neuroscience and machine learning into closed-loop experiments.

## Introduction

In neuroscience and neuroengineering, researchers use closed-loop systems to respond to neural activity in real-time – for example, to stimulate a neural circuit or control an end effector – in order to test properties of the brain or build devices that interface with it for a therapeutic purpose. In research with intracortical BCIs (iBCI), closed-loop systems have enabled people with paralysis to control a robotic arm, spell sentences, move limbs with functional electrical stimulation, and speak [1]–[6].

Closed-loop systems must meet stringent timing requirements that are derived from the timescales of the neural processes being studied. Neuronal action potentials, or “spikes”, have a waveform on the order of a millisecond, so systems that process spiking data need to acquire the waveform at a sub-millisecond resolution and detect spike events at 1 kHz resolution. Another important factor is the control latency, i.e., the time it takes to produce a control signal from a window of neural activity. For applications such as using an iBCI to control a computer cursor, the latency from neural recording to behavioral prediction has typically been 15-20 ms [3], [7]; increasing latency is known to decrease control performance [8], [9]. We thus want a system that can receive and process spiking data at 1 kHz and predict movement intention with less than 15 milliseconds of latency.

Several groups have released software packages for building real-time systems, but they lack the features needed to run both the highly-optimized C/C++ code that handles data acquisition and the modular Python code that modern machine learning libraries are built in. They are typically designed to run code in a limited set of programming languages, making it difficult to find a system that provides broad support for the many high-level languages used in the neuroscience field, including Python, MATLAB, and Julia. Simulink Real-Time (Mathworks) provides a visual programming interface that supports real-time MATLAB and C/C++ code. RTXI and Falcon provide sub-millisecond timing guarantees but are restricted to running C/C++ code [10], [11]. LiCoRICE supports interprocess communication between Python and C, but it does so via Linux shared memory primitives (i.e., semaphores and shared memory objects), and hence has a high barrier-to-entry for most neuroscientists (**Table S1**). These existing systems lack the language support and communication mechanisms needed to run ANNs natively in their original programming environment, presenting a barrier to deploying ANN models in closed-loop experiments.

Partially due to the challenge of deploying them in real-time, computational models that are promising for closed-loop neuroscience applications often go untested in a closed-loop setting. These models were developed with offline analyses on existing data, which are important for rapidly iterating on model architectures and hyperparameters but fall short of capturing the real-time response of the brain in situations involving closed-loop feedback. For example, closed-loop feedback is known to be a critical component of iBCI control, so, in the iBCI context, offline analysis can provide only a partial validation of a decoding model [8], [12]. Evaluating such models in closed-loop iBCI experiments would be a worthwhile research direction, but the space of models that can be tested is limited by the labor-intensive (and potentially error-prone) process of reimplementing models that contain millions of parameters to achieve compatibility with existing closed-loop software architectures. Thus, there is a “translation gap” between the development of new computational models and the evaluation of those models in closed-loop experiments.

We developed the Backend for Realtime Asynchronous Neural Decoding (BRAND) to address three critical needs: (1) running closed-loop ANN inference in the same runtime environments used for offline analyses, (2) supporting several programming languages, and (3) providing sub-millisecond, high-bandwidth communication between system processes. This system uses the popular Redis in-memory database, which provides it with low-latency IPC and broad compatibility across the 54 different programming languages for which Redis client libraries exist [13]. With BRAND, researchers can easily integrate a variety of computational models into their experimental pipelines by structuring those models to read from and write to the Redis database. Computational models can then integrate with the system components responsible for signal processing, behavioral tasks, and visual feedback with sub-millisecond communication latency. This results in a system that makes it easier to prototype and study new computational models, processing techniques, and behavioral tasks in a research setting.

In this paper, we validate BRAND in three contexts: (1) high-bandwidth interprocess communication, (2) BCI control with ANNs, and (3) neural data simulation. BRAND’s interprocess communication latency was consistently less than 600 microseconds with inputs of up to 1024 channels of 30 kHz neural data. In the BCI control benchmark, BRAND runs all of the components needed to acquire and process 30 kHz neural data over a network and produce hand movement predictions with an ANN in less than 8 milliseconds. Using this pipeline, we conducted a closed-loop demonstration of iBCI cursor control with a participant in the BrainGate2 clinical trial (CAUTION: Investigational device. Limited by federal law to investigational use.). BRAND was also used to generate simulated neural data for both speech decoding and cursor control. With these results, BRAND is shown to be a versatile system for building a wide range of closed-loop experiments with low latencies and full support for state-of-the-art machine learning techniques.

## Methods

### System Architecture

BRAND follows a modular design that divides the control of a neuroscience experiment into small reusable components, each of which is designed to complete a part of the overall computational pipeline. We refer to each component as a *node*; several nodes are combined together to form a *graph*. For example, a minimal graph for an iBCI experiment might consist of a feature extraction node (e.g., computing local field potential power or spike rate), a decoding node, and an effector control node (**Fig. 1a**). Nodes run in parallel as separate processes, which allows us to decouple critical, highly-optimized code like data acquisition from slower, experimental code like neural network inference. Data are passed between nodes asynchronously via *Streams* in a Redis database. Each node can publish data to several output streams, and receive data by subscribing to several input streams. These streams provide a straightforward way to build a chain of interchangeable nodes that run in parallel. Running nodes in parallel allows BRAND to process incoming data at a higher rate than an equivalent system that runs sequentially (**Fig. 1b-c**). Streams are append-only logs that persist in the database even when data have already been read, so they also maintain a record of all data within an experiment and can be saved for later analysis.

**Figure 1.**
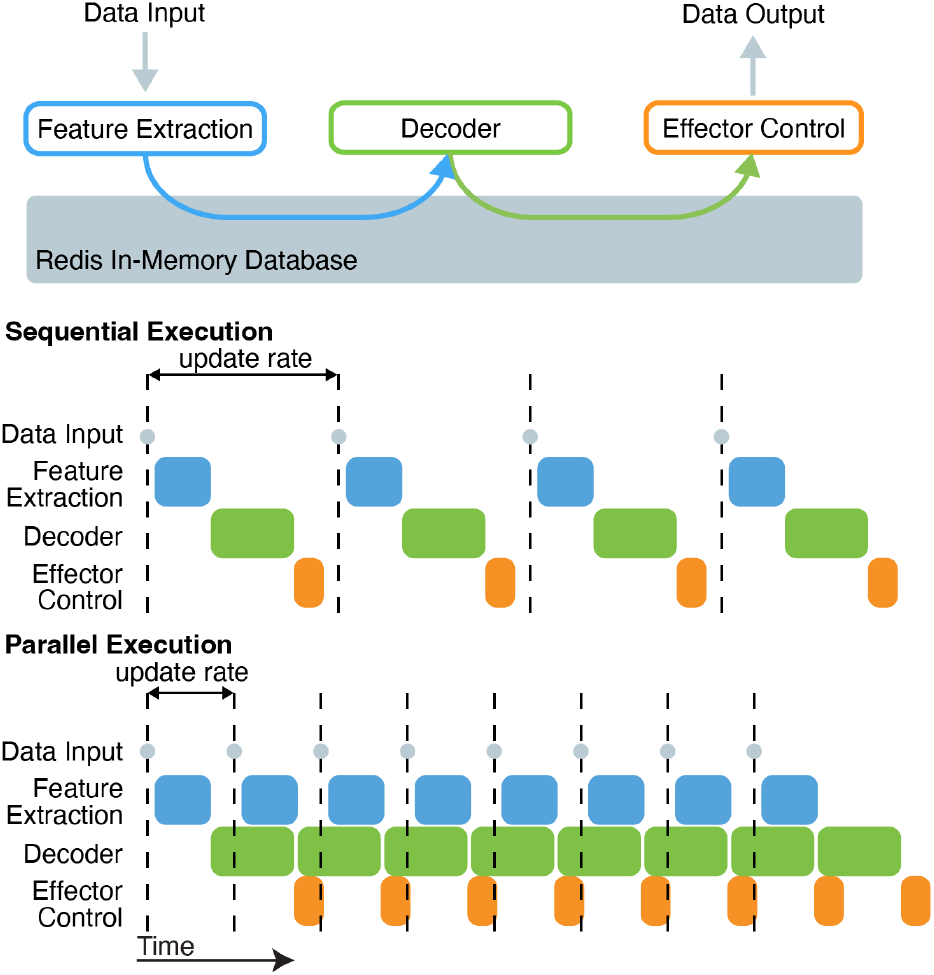
Software architecture schematic. a) BRAND consists of a set of processes, or “nodes’’, that each receive inputs and/or produce outputs during an experiment. b) If nodes were run sequentially (as if they are in a script), all nodes would need to finish processing a given sample before the next one could be processed. Delays in any part of the processing chain would cause the whole system to fall behind and delay critical events like acquiring an incoming sample. c) In BRAND, nodes run in parallel and communicate asynchronously, allowing them to maximize the rate at which data are processed and minimize the chance that delays in downstream nodes would cause the system to fall behind.

In BRAND, graphs for closed-loop experiments are configured using Yet Another Markup Language (YAML). Each graph is configured by a single YAML file that lists the nodes that will be run in the graph and the parameters for each of those nodes. A script, called *supervisor*, parses this YAML file and then initializes the specified nodes as separate Linux processes. The parameters for the graph are then sent to those nodes through Redis. Once a graph is loaded, the supervisor may then start or stop nodes according to commands received via Redis. In a typical experimental session, a researcher will supply a list of graphs (as YAML files) that each configure the system for a different step of the experiment. The researcher will start and stop these graphs by sending commands to supervisor. BRAND provides flexibility in this experimental workflow, allowing researchers to control the system with the Redis command line or develop their own graphical user interfaces for selecting graphs and monitoring or adjusting the parameters of their nodes. Since Redis enables communication across multiple computers on the same network, BRAND includes another launcher script (called “booter”) that extends *supervisor’s* capabilities to additional machines. To run nodes across several machines, the researcher would start a *supervisor* instance on the machine that hosts the Redis database and start a *booter* instance on each client machine. Nodes can then be configured to run on any of these machines and send and receive data from the common Redis database. Users can also configure the process priority and processor affinity for each node to make full use of the resource allocation tools available in Linux.

To maintain access to the full suite of Redis features, BRAND does not impose any requirements on the way in which programmers interface with the Redis database. However, programmers are encouraged to use the Stream data type for communication between nodes, as it facilitates data logging, and provides a standard asynchronous communication mechanism between nodes. Libraries for Redis exist in many different languages, including C, Python, Go, Julia, and MATLAB [13]. Redis can either be configured for communication via Transmission Control Protocol (TCP) sockets, which allows multi-computer communication, or Unix sockets, which allows for faster interprocess communication within a single machine.

In summary, BRAND was designed with a modular node-graph structure that is configurable via YAML files and uses Redis for communication across multiple programming languages and multiple computers. With these design choices, we aim to make it easy to both integrate and share individual components of an experimental pipeline across experiments and labs.

### Validation

We tested BRAND on the Linux operating system. Most demonstrations in this manuscript were conducted using Ubuntu 20.04 LTS and the Linux kernel with a PREEMPT_RT patch (version 5.15.43-rt45) [14] and run on a Dell Optiplex 7000 small form factor PC with an Intel i9-12900 processor and 128 GB of memory. The speech simulator benchmark was conducted on an AMD Ryzen 9 7950X processor running Ubuntu 22.04.2 LTS and Linux kernel 5.19.0-41-generic (without the PREEMPT_RT patch).

#### Communication Latency

To test the communication latency of the Redis database, we ran a benchmark in which packets approximately matching the size of 30 kHz neural data from the Blackrock Neural Signal Processor (Blackrock Neurotech, Salt Lake City, Utah, USA) (NSP) were sent from a *publisher* node to a *subscriber* node. We recorded timestamps from the system clock (with Python 3.8.2’s *time*.*monotonic_ns()* function) before the data were written to Redis in the publisher and after the data were read from Redis in the subscriber. The difference between these two timestamps was considered the “communication latency” of the Redis database when using TCP sockets on a single machine. For each test condition we measured 300,000 pairs of timestamps. Both the publisher and subscriber nodes were implemented in Python and then compiled using the Cython package (version 0.29.18).

We ran three variations of this benchmark when varying (1) the number of input data channels, (2) the publishing rate, and (3) the number of nodes. In all tests, publisher packets were encoded as arrays of 16-bit integers, and the number of values in each array was 30 times the number of channels (since we sent one packet of 30 kHz data every millisecond). In the first test, a publisher node sent packets at 1000 Hz and a subscriber node read each of them and logged the timestamp at which it received them. This test was repeated for four different packet sizes: 128, 256, 512, and 1024 channels. In the second test, the number of channels was held constant at 128, and the output rate was set at 100, 200, 500, or 1000 Hz. In the third test, 2-4 nodes were chained together and all but the last node forwarded the data they had received as input. This test was run with 128 channels and a 1000 Hz sampling rate (**Fig. 2a**).

**Figure 2.**
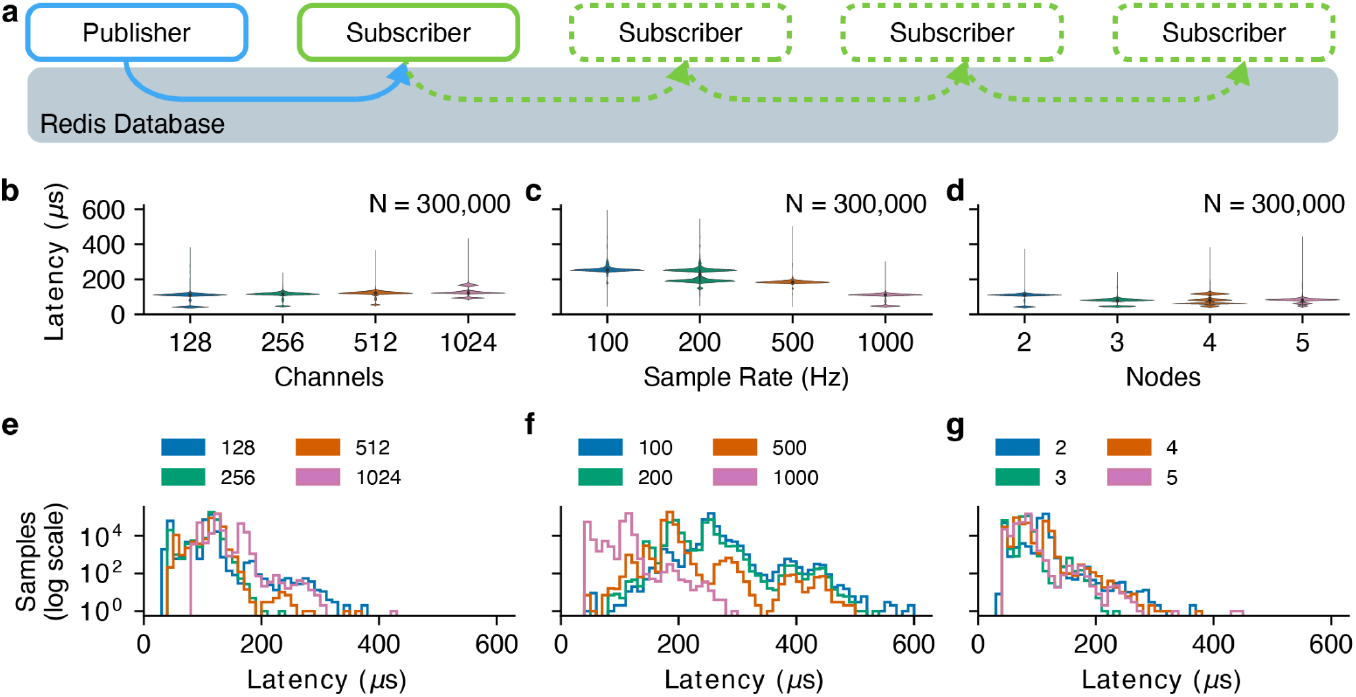
BRAND achieves low-latency inter-node communication. a) To test inter-node communication latency, a publisher node sends 30 kHz neural data (grouped in 1-millisecond packets) to a subscriber node via the Redis database. Violin plot of the resulting latency measurements showing that the inter-node communication latency is consistently below 600 microseconds even as b) the channel count is scaled up to 1024 channels, c) the sampling rate is changed, and d) additional subscriber nodes are added. Histograms of these data show the distribution of latency measurements for each e) channel count, f) output rate, and d) number of nodes.

#### iBCI Control

To evaluate the latency of BRAND when performing the processing tasks needed for an iBCI, we benchmarked an iBCI control graph with two different decoders. In this benchmark, simulated 30 kHz data from two NSPs (firmware version 6.05.02) with a Neural Signal Simulator (Blackrock Neurotech, Salt Lake City, Utah, USA) were acquired over the network and then filtered with a 250 Hz high-pass filter applied forward and backwards to a 4 ms buffer of the incoming data. The filtered data were then thresholded at −3.5 times the root-mean-square voltage of each channel to detect threshold crossings within each 1 ms window. These threshold crossings were then binned into 10 ms bins and normalized before being passed to a decoder. The decoder’s cursor velocity predictions were smoothed with an exponential moving average and scaled before the cursor position and task state were updated (**Fig 3a, Fig S1a**) [15], [16].

**Figure 3.**
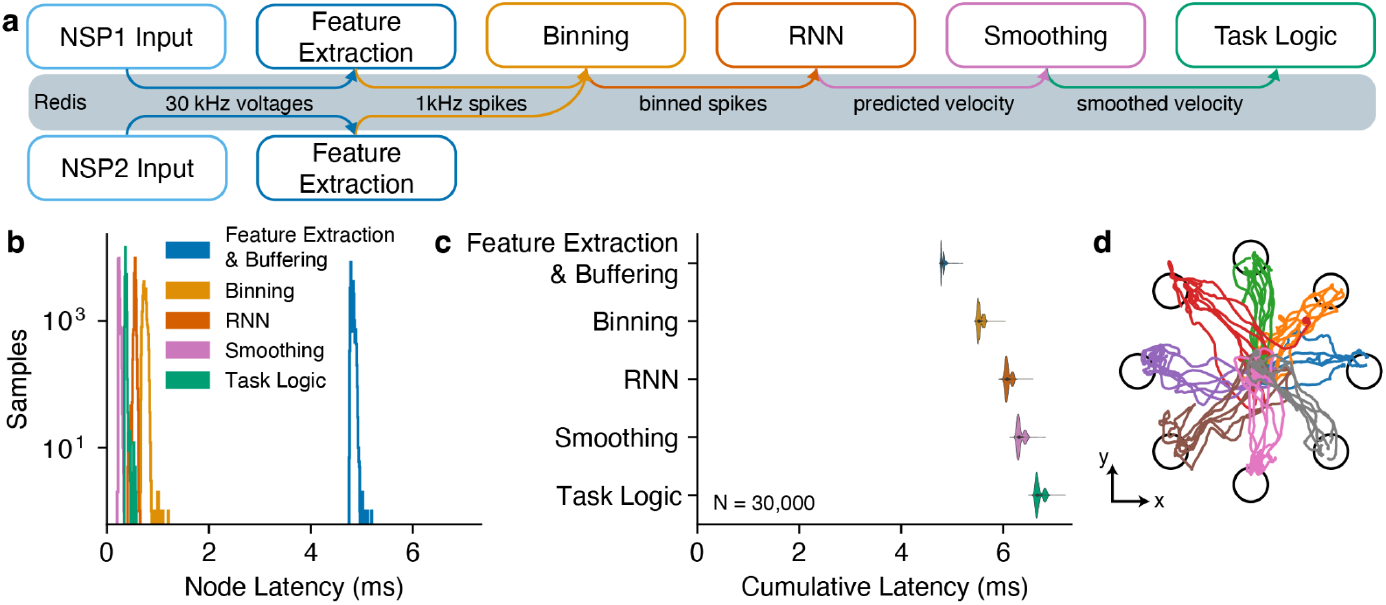
BRAND can be used for low-latency iBCI control. a) To test end-to-end iBCI control latency, we ran a graph that received 30 kHz 96-channel neural spiking data via UDP (Ethernet) from two Blackrock NSPs (total of 192 channels), extracted spiking features at 1 kHz, binned spikes into 10-millisecond bins, ran decoding, and updated the location of the cursor in the task. This test used a recurrent neural network (RNN) decoder. This graph was benchmarked using simulated data. b) Latency measurements for each node were plotted as histograms (N = 30,000 packets). c) The cumulative latency is plotted relative to the time at which each node (vertical axis) wrote its output to the Redis database. On the horizontal axis, zero is the time at which the last sample in each bin was received over the network from the NSPs. d) Cursor positions during iBCI-enabled cursor control.

For neural decoding, we implemented two different decoding algorithms: an optimal linear estimator (OLE) and a recurrent neural network (RNN). The OLE decoder estimated the cursor velocity as a linear combination of the input features at a single time step. The OLE decoder had a 384-dimensional input consisting of threshold crossings and spike-band power [17] from 192 channels of neural data. Spike-band power was computed by applying a 250 Hz high-pass filter to the 30 kHz neural data, squaring the result, and averaging across time within each 1 ms window. Each feature was normalized using its mean and standard deviation from the previous period of neural recording. This decoder was fit using ridge regression, with the weight of the L2 regularization term being chosen via 3-fold cross-validation. The RNN decoder was implemented in PyTorch 1.12.1 and PyTorch Lightning 1.7.1 and consisted of a 76-unit long short-term memory (LSTM) layer followed by a 2-unit linear fully-connected layer. This decoder received 192 channels of binned threshold crossings as input and applied normalization using the mean and standard deviation of the training data. Its L2 regularization, dropout, learning rate, and LSTM dimensionality were chosen using a random search on previously-collected data. During the session, this decoder was trained on four minutes of closed-loop cursor control data with the OLE decoder using an Nvidia GeForce RTX 3090 graphics processing unit (GPU).

To highlight the capabilities of the BRAND system, we also ran real-time inference with two large neural networks designed for feature extraction: the Neural Data Transformer (NDT) and Latent Factor Analysis via Dynamical Systems (LFADS). These autoencoder networks denoise neural data by modeling the dynamics of population-level activity and could be useful additions to many closed-loop neuroscience experiments [18], [19]. Both networks were run as Cython-compiled nodes in BRAND using their published PyTorch (version 1.12.1) and Tensorflow (version 2.2) implementations. Inference latencies were measured by comparing the monotonic clock timestamps between each node’s input and output.

#### Simulation

We also benchmarked the timing performance of BRAND when operating as a neural data simulator. For this, we implemented two different simulators: one for cursor control and one for speech decoding. In the cursor control simulator, the human-controlled movements of a computer mouse were translated into firing rates via a cosine tuning model. In the speech simulator, audio spoken into a microphone was encoded as mel-frequency cepstral coefficients (MFCCs) [20], which were then used to generate neuronal firing rates. Both simulators then generated 96 channels of 30 kHz simulated voltages from the firing rates and broadcast them using the same packet structure as Blackrock NSPs. Latency measurements were taken using timestamps logged immediately prior to writing the output of each simulator node to Redis.

### Closed loop validation

#### Clinical Trial Participant

T11 is an ambidextrous man, 38 years old at the time of the study, who had suffered a C4 AIS-B spinal cord injury approximately 11 years prior to enrolling in the BrainGate2 clinical trial (ClinicalTrials.gov Identifier: NCT00912041). Neural data were recorded from two 96-channel microelectrode arrays placed in the “hand knob” area of his left (dominant) dorsal precentral gyrus in the BrainGate2 trial. Data were collected on Trial Day 1321 (1,321 days after surgical placement of the arrays). The Institutional Review Boards of Mass General Brigham (#2009P000505) and Providence VA Medical Center granted permission for this study.

#### Cursor Control Task

We asked participant T11 to perform the classic radial-8 center-out-and-back cursor control task [21]. This experiment was organized into “blocks”, which are periods of data collection lasting 3-4 minutes. During these experiments, T11 was seated comfortably in his wheelchair and looked at a computer monitor. We began with an open-loop cursor calibration block, in which the cursor automatically moved to targets which T11 was asked to attempt to follow with his right thumb as if using a joystick. We then trained an OLE decoder to predict the cursor’s velocity from neural data [22]. In the next block, we set up the OLE decoder to control the cursor velocity and asked him to attempt to move it to the active target. Data from this block were used to train a recurrent neural network (RNN) decoder to predict his intended cursor velocity. The intended cursor velocity was approximated as the vector resulting from subtracting the cursor’s position from the target’s position scaled to the original OLE prediction’s magnitude or zeroed if the cursor reached the target [23]. In a later block, T11 was asked to perform the radial-8 task again with this RNN decoder. The initial open-loop block lasted three minutes while each of the subsequent closed-loop blocks lasted four minutes.

#### Signal Processing

Neural data were filtered by the Blackrock NSP at 0.3 Hz to 7.5 kHz and then broadcast in UDP packets over Ethernet. These data were acquired and published to Redis with a BRAND node and then, in other nodes, the data were common-average referenced, filtered, and thresholded to yield threshold crossing and spike-band power features. Those features were then grouped into 10 ms bins, normalized, and sent into the decoder node [7]. The decoder’s predictions were smoothed exponentially and scaled according to a gain parameter and then passed to a finite-state machine (FSM) node that updated the cursor’s position and task state. Finally, a graphics node rendered and displayed the task on a separate PC according to the cursor and target information sent by the FSM.

## Results

### Inter-process communication latency

We evaluated BRAND using a publisher-subscriber benchmark, where 30 kHz neural data emitted at 1 kHz (chosen to reflect the sampling rates of the Blackrock NSP system for intracranial recordings) were streamed from the publisher to the subscriber in one-millisecond packets. In this test, we found that BRAND had a communication latency of less than 600 microseconds when sending up to 1024 channels of neural data. This held true for several sampling rates and also when we increased the number of subscribers in the graph from one to four (**Fig. 2**). These results indicate that BRAND’s chosen communication mechanism, Redis, is fast enough to consistently transmit the high-bandwidth data encountered in neural recordings with sub-millisecond latency.

### iBCI Control

To evaluate the practical application of BRAND in an iBCI control setting, we developed a benchmark in which 30 kHz neural data were acquired, filtered, and thresholded to obtain spiking features (see Methods for details). The features were then binned and passed into two different types of decoders (OLE and RNN), and the decoder predictions were sent to a node that controls the task state.

For these decoders to be considered real-time, they needed to process incoming data within a set latency deadline and with minimal jitter. In this case, for a 10 ms bin size, a new sample arrives at the decoder every 10 ms. Real-time processing would be achieved by having a per-node latency of less than 10 ms for each of these decoders. When using the OLE decoder, we found decoding latencies to be consistently less than 0.6 ms per 10 ms bin decoding step (**Fig. S1**). With the RNN decoder latencies were consistently below 1.2 milliseconds per step (**Fig. 3**). Both of these decoders were well-below the 10 ms-per-step deadline for real-time processing.

In closed-loop experiments with these decoders, participant T11 performed a task in which he moved a cursor to one of eight radially-arranged targets on a screen and then returned the cursor to the center. T11 achieved a median target acquisition time of 1.76 ± 0.63 seconds with the OLE decoder and 1.79 ± 0.74 seconds with the RNN decoder. This performance was consistent with previous demonstrations of this now-standard cursor control BCI task [7], [24].

To further evaluate the advanced ANN inference capabilities that BRAND provides, we sought to benchmark the latencies of two state-of-the-art latent variable models: Latent Factor Analysis via Dynamical Systems (LFADS) and Neural Data Transformer (NDT) [18], [19]. LFADS and NDT were designed to improve the extraction of neural features for use in decoding, and have previously achieved a velocity prediction R^2^ of 0.9097 and 0.8862, respectively in the “MC_Maze 5 ms” dataset in the Neural Latents Benchmark [25], which is a marked improvement over the 0.6238 achieved using smoothed binned spikes. Each model was made up of several neural network layers that rely on specialized Python libraries (Tensorflow and PyTorch) for training and inference. Previous real-time systems that lacked Python support would have required a full reimplementation of these models in another language like C or C++ to run this test [11], [26]. With BRAND, we can simply connect the existing TensorFlow and PyTorch implementations of these models to the fast Redis IPC mechanism that is used throughout the system. In our testing, LFADS inference times were consistently below 6 ms and NDT inference times are below 2 ms, with both models staying under our 10 ms latency criterion for real-time inference (**Fig. 4**).

**Figure 4.**
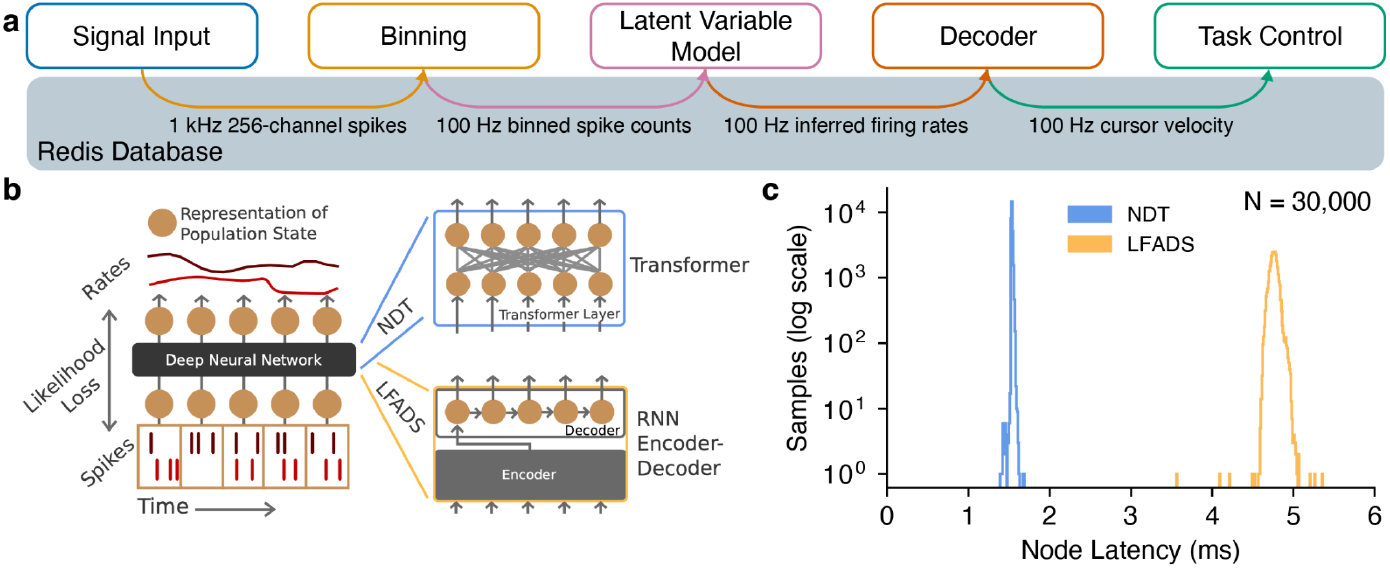
BRAND runs ANN latent variable models with low latency. a) To test the inference latency of LFADS and NDT, we inserted them into an iBCI control graph that receives 256 channels of simulated threshold crossings at 1 kHz, bins them, runs inference with LFADS or NDT, runs decoding, and updates the task state. b) LFADS and NDT use different types of sequence models, an RNN and a Transformer, respectively. Reprinted from Ye and Pandarinath 2021 [18]. c) NDT inference times are consistently below 2 ms, while LFADS inference times are consistently below 6 ms.

### Neural Simulation

Neural data simulators have been a critical component of neuroscience research, particularly in the clinical iBCI research. Simulators enable the testing of iBCI decoding algorithms and tasks before they are used in a clinical research setting, where time with participants is extremely limited and valuable. This allows researchers to rigorously validate their signal processing, decoders, and tasks prior to an experiment. Full system simulation also helps lower the chance that software issues will impede data collection.

We evaluated whether BRAND was able to act as a real-time simulator for testing iBCI applications. With its modular design, we can use interchangeable inputs and firing rate encoding models to support simulating neural activity during different types of behavior. In this case, we tested two simulators: one for cursor control and one for a speech decoding. We demonstrated that BRAND can run both simulations in real-time. The cursor control simulator runs with less than 4 ms of end-to-end latency and the speech simulator runs in less than 3 ms (**Fig. 5**).

**Figure 5.**
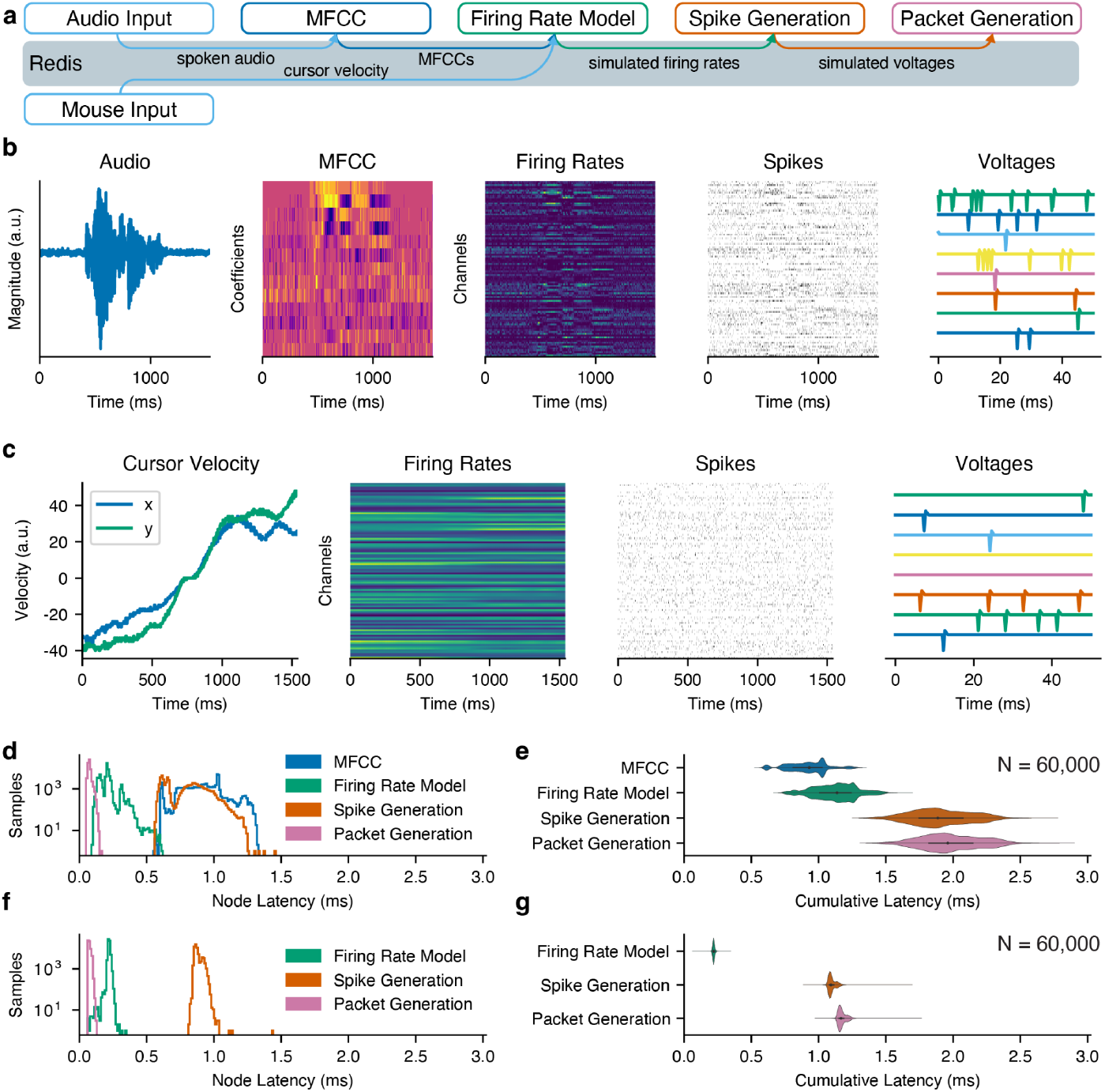
BRAND enables low-latency, real-time simulation of neural data. a) In the cursor control simulator, computer mouse movements are translated into neural firing rates with a cosine tuning model and then used to generate and broadcast 30 kHz voltage recordings as ethernet packets. In the speech decoding simulator, spoken audio is used to generate neural data based on simulated neurons with weights tuned randomly to different spectral features. b) Examples of data recorded from the cursor control simulator. c) Examples of data recorded from the speech simulator. d) Latency of each node in the speech simulator and e) cumulative latency. f) Latency of each node in the cursor control simulator and g) cumulative latency.

## Discussion

We introduce and validate the BRAND Real-time Asynchronous Neural Decoding system, a real-time software platform that aims to meet the need in contemporary experimental neuroscience and neuroengineering for software frameworks that support both ANN inference in Python and low-latency control for intracortical BCIs. BRAND fills this need by providing three critical features: (1) running closed-loop ANN inference in the same runtime environments used for offline analyses, (2) supporting a wide range of programming languages, and (3) providing sub-millisecond high-bandwidth communication between system processes. The latency and jitter of the system were benchmarked in three settings: streaming of high-bandwidth neural data, simulation of neural data from user input, and iBCI control of a computer cursor. We demonstrated that BRAND can perform low-latency data acquisition in C and Python while also running real-time inference with multiple neural network libraries. By providing a language-agnostic framework for real-time software, BRAND reduces the need to rewrite complex computational models when integrating them into a real-time experiment, allowing researchers to more rapidly explore and validate novel computational models.

### Open-Source Software

The BRAND code is publicly available, and users are encouraged to develop plugins (nodes) that make BRAND work with additional recording systems, models, and tasks. The open-source approach provides two main benefits: code duplication is minimized and replicating studies across research groups is easier. BRAND’s dependencies, including Python, Redis, and Linux, are also free and open-source, so users can build experimental systems without having to pay software licensing fees. The current source code is at: github.com/brandbci/brand. The code has been released under the MIT license.

BRAND’s modular design supports seamless code sharing across labs. By standardizing around a common IPC mechanism with Redis, individual nodes (like models or signal processing steps) can be integrated into experimental pipelines in different labs without the need to restructure other components of each lab’s systems. For example, two labs with different data acquisition systems can use the same decoding models and behavioral tasks while maintaining separate implementations of their data acquisition node. Similarly, a computational lab can develop and share a real-time implementation of their new decoding model, and labs that conduct experiments can integrate that model into their experimental pipeline without needing to modify their existing data acquisition code. Developers in the wider neurotech industry have also begun to develop tools that expand the capabilities of BRAND by adding features like real-time data visualization (**Fig. S2**).

### Comparison to Existing Systems

Given the fundamental importance of software in performing high-quality neuroscience experiments, several groups have released software packages for closed-loop experiments. Systems like Simulink Real-Time [7], RTXI [10], and Falcon [11] provide tight timing guarantees but lack the Python support that is essential for machine learning research. Timeflux [27] and LabGraph [28] solely support Python, and thus suffer from the lack of support for lower-level languages like C that are more suitable for latency-critical applications. LiCoRICE pairs Python and C support, but does so via shared memory primitives that are challenging to use and lack support for multi-machine communication [29].

These competing systems all require developers to choose from a limited set of programming languages, which restricts the pool of existing libraries that can be used when developing experiments. For example, the machine learning community has standardized around the use of Python, and several libraries (e.g., PyTorch, TensorFlow, Keras, JAX) have been developed to perform the difficult task of training artificial neural networks on GPUs and other hardware accelerators. Meanwhile, the C programming language has long been the foundation of Linux development and offers the low-level control of memory management that is often critical for optimizing compute latency, as well as interfacing with input/output devices and other peripherals. Other languages like Julia, R, and MATLAB are widely used for computational research. Javascript and its derivatives play an important role in web development and GUIs. C# and Java are widely used in game (and thus task) development. Dareplane, another recently-published real-time framework has been designed to provide broad language compatibility, but unlike BRAND, it has not been validated to support the high-bandwidth communication needed for iBCIs that contain hundreds of channels [30].

### Distributed Computing

It is often useful to distribute computational tasks like signal processing, model training, and graphics rendering across multiple machines to make use of additional compute resources and avoid overloading a single computer. For this to work in a real-time system, processes running on each machine need to be able to pass data between one another with low latency. BRAND readily supports distributed computing, since Redis is designed to work across large clusters of computers that rapidly share data and coordinate to perform large-scale computational tasks. Communication with the Redis database is done with the same commands and syntax whether the database is hosted locally or remotely, so nodes can seamlessly be moved from machine to machine without changing their implementation. We used this feature extensively in our development to run simulators and GPU-related code (such as neural network training and graphics rendering) on their own dedicated machines to avoid interfering with the latency-critical signal processing that occurs on our real-time decoding machine.

### Future Directions

While the experiments shown in this study were conducted on stereotyped block design-based tasks, BRAND could potentially be extended to personal use settings [31], where the participant would use an iBCI for their daily activities. BRAND’s modular structure provides a straightforward path to making nodes that can be hot-swapped without interrupting device use. For example, a participant may want to calibrate and deploy a new decoder every few hours to maintain high-performance control or select a different decoder for different tasks. BRAND could serve as a useful environment for prototyping this kind of personal use software in research studies.

BRAND also fills a critical need in speech iBCI research, where artificial neural networks have enabled several recent advances in brain-to-text decoding [6], [32] and are anticipated to play an important role in the development of real-time speech synthesis BCIs [33]. Unlike text decoding, speech synthesis is expected to benefit from millisecond-scale closed-loop feedback that mimics the way in which able-bodied people can hear their own voice while speaking. BRAND is uniquely suited to this research area by providing the combination of ANN support and sub-millisecond IPC latency that is needed for a real-time speech synthesis BCI.

BRAND could also become a useful tool for neurological research in general. The development of new computational models is a critical avenue for studying neural activity across several brain functions, including movement, sensation, and cognition [34], [35]. Advances in modeling and decoding neural activity could lead to therapeutic benefits if paired with devices that interface with the brain, like responsive neurostimulators for epilepsy [36]. Existing clinical systems for responsive neuromodulation or deep brain stimulation allow streaming of data over Ethernet and could possibly be integrated into BRAND for use in research studies.

## Acknowledgments

The authors would like to thank Participant T11, his family and caretakers, Beth Travers, Dave Rosler, and Maryam Masood for their contributions to this research. We also thank Antonio Eudes Lima, Diogo Schwerz de Lucena, and Robert Luke at AE Studio for their development of the Neural Data Visualizer. This work was supported by the Emory Neuromodulation and Technology Innovation Center (ENTICe), NIH Eunice Kennedy Shriver NICHD K12HD073945, NIH-NINDS/OD DP2NS127291 (CP), NIH-NIDCD/OD DP2DC021055, Simons Collaborations for the Global Brain Pilot Award 872146SPI (SDS), NIH NIBIB T32EB025816 (YHA), NIH F32HD112173 (SRN), NIH-NIDCD U01DC017844, and Department of Veterans Affairs Rehabilitation Research and Development Service A2295R and N2864C (LRH). The content is solely the responsibility of the authors and does not necessarily represent the official views of the National Institutes of Health, or the Department of Veterans Affairs, or the United States Government.

## Data availability statement

All data used to produce the figures in this paper will be available upon publication.

## Author contributions

YHA, DMB, and CP conceived the project. YHA, KB, MR, KP, NSC, BB, SRN, DMM, XH, and DMB contributed to software design and development for BRAND. YHA developed and ran benchmarks to validate the system and wrote the manuscript. MR and NSC developed and benchmarked the neural data simulators. CN, YHA, SRN, MR, and DMM conducted the experiments with participant T11. SA and SRN deployed the software and hardware needed for the T11 experiments. LRH is the sponsor-investigator of the multi-site clinical trial. DMB, CP, LEM, SDS, and NAY supervised and guided the project. Funding was acquired by CP, DMB, SDS, and LRH. All authors reviewed and contributed to the manuscript.

## Supplemental Material

**Figure S1.**
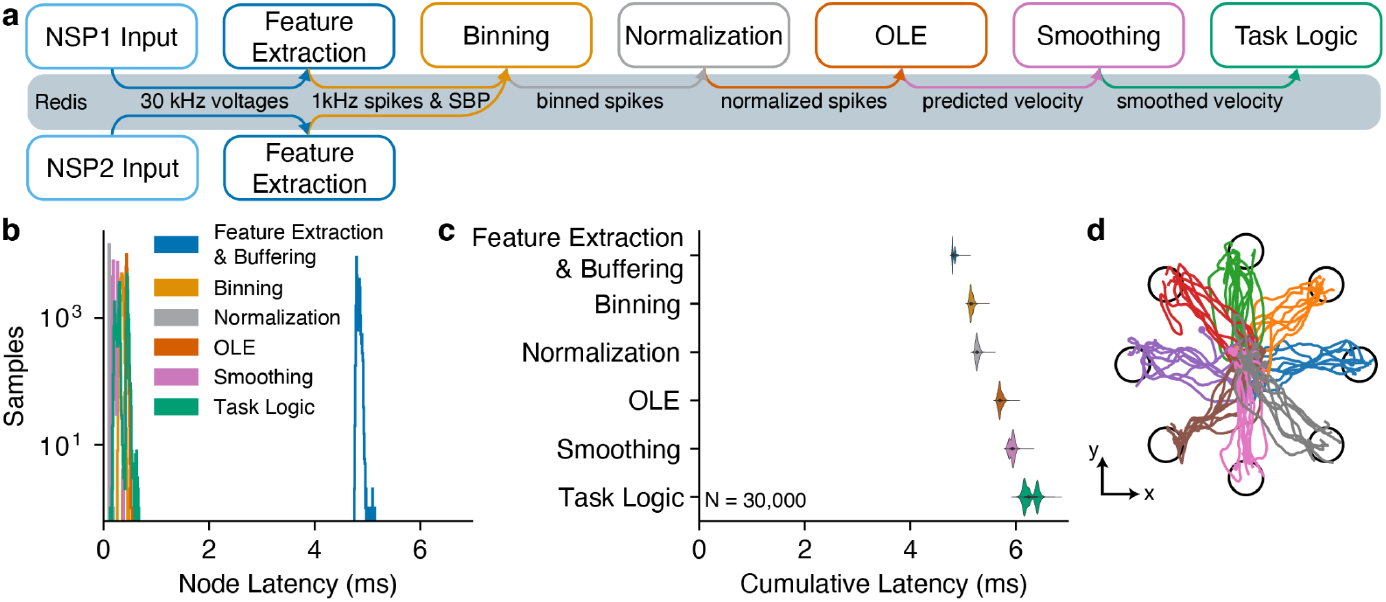
iBCI Control with an OLE decoder. a) To test end-to-end iBCI control latency, we ran a graph that receives 30 kHz 96-channel neural spiking data via UDP (Ethernet) from two Blackrock NSPs (total of 192 channels), extracts spiking features at 1 kHz, bins spikes into 10-millisecond bins, runs decoding, and updates the location of the cursor in the task. This test used an OLE decoder. b) Latency measurements for each node are plotted as histograms (N = 30,000 packets). c) The cumulative latency is plotted relative to the time at which each node (vertical axis) writes its output to the Redis database. On the horizontal axis, zero is the time at which the last sample in each bin was received. d) Cursor positions during closed-loop control.

**Figure S2.**
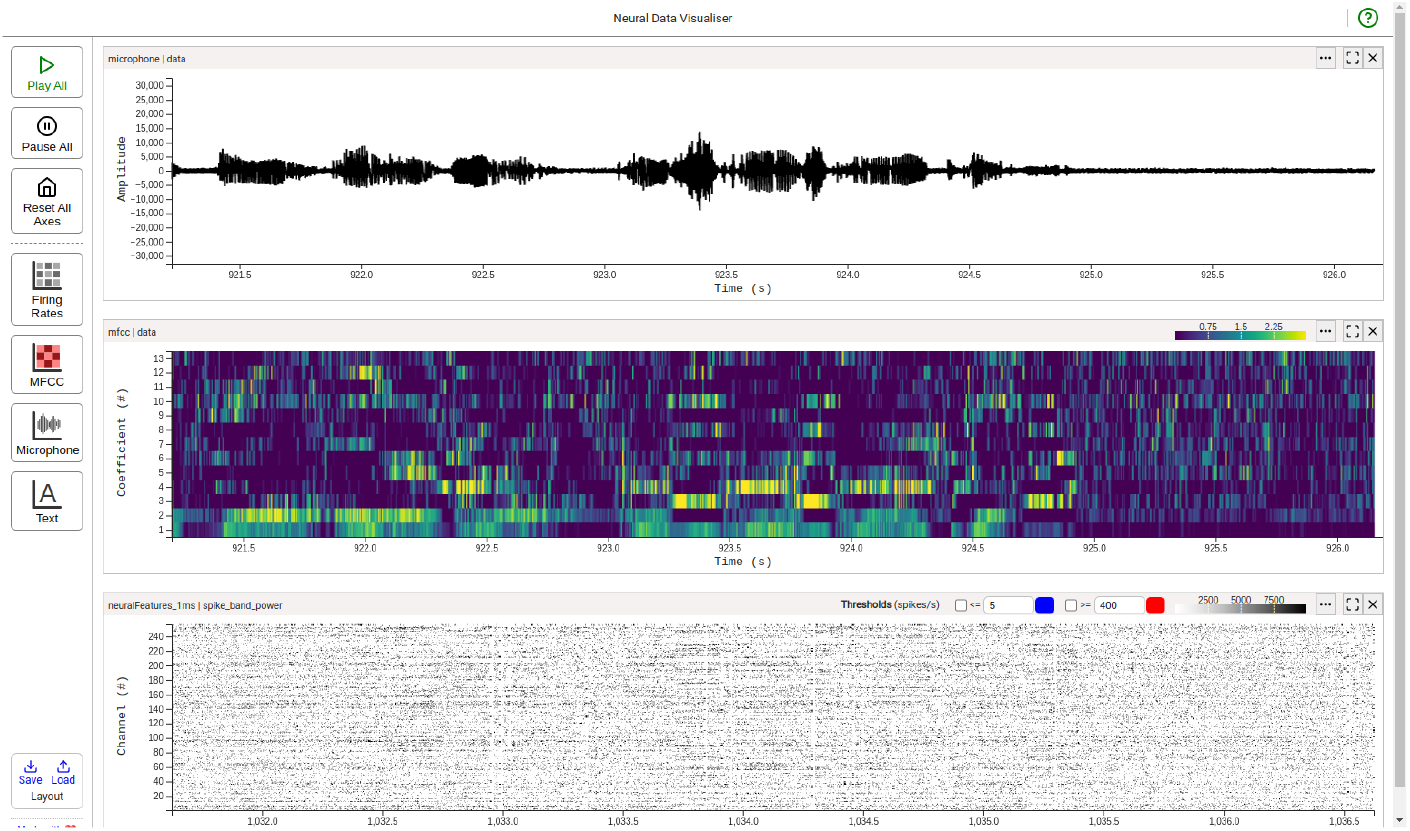
Neural Data Visualizer enables real-time data visualization. This real-time visualization tool developed by AE Studio (Los Angeles, USA) allows users to visualize data in their experiment’s Redis Streams in real-time. This visualizer was developed independently of BRAND with only very lightweight coordination (the authors provided an example of their BCI systems’ Redis Streams) and serves as an example of the add-ons that can be used to expand BRAND’s capabilities.

**Table S1.**
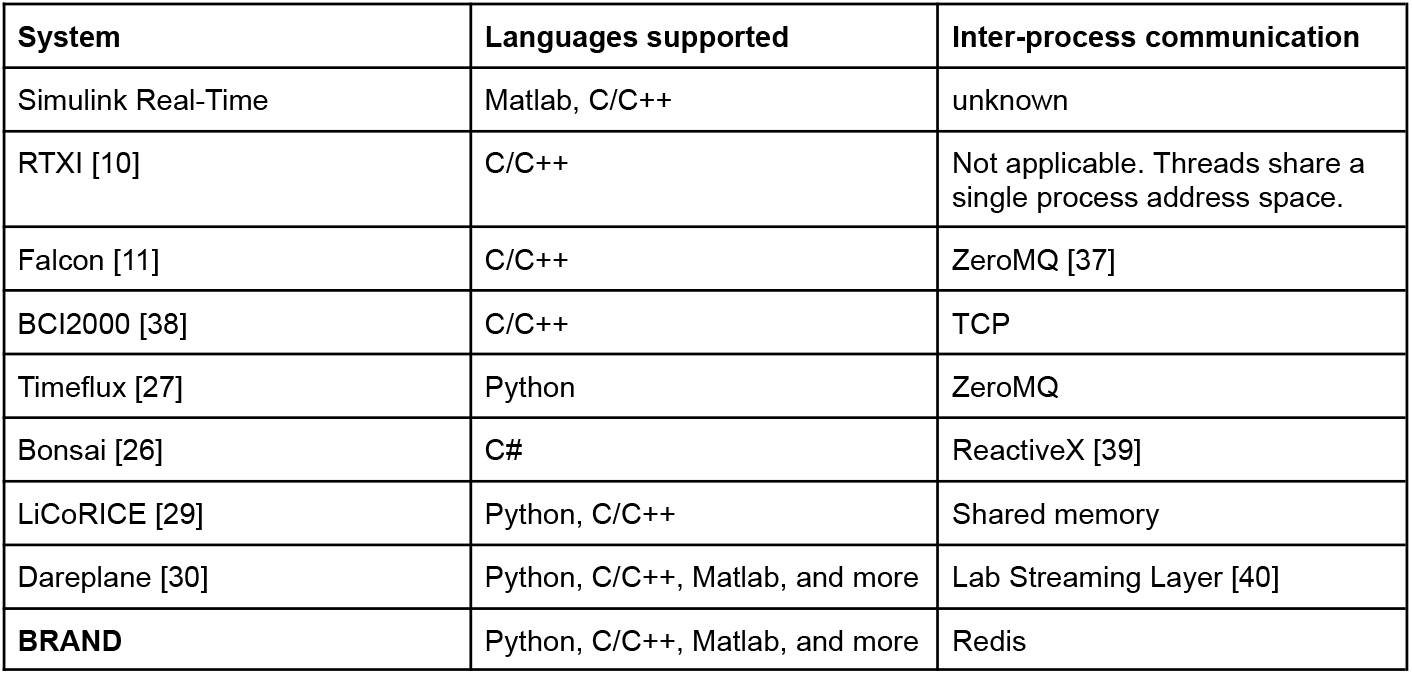
Real-time systems in neuroscience.

